# Label-free Isolation of Heterogeneous Breast Cancer Cell Populations via Insulator-Based Dielectrophoresis^†^

**DOI:** 10.64898/2026.08.04.742571

**Authors:** Gürhan Özkayar, Ismene N. Usman, Ece Yakın, Jaco Kraan, Kristen David, Duco Bosma, John W.M. Martens, Peter ten Dijke, Georg R. Pesch, Pouyan E. Boukany

## Abstract

Circulating tumor cells (CTCs) are valuable biomarkers for cancer diagnosis and monitoring, yet their isolation from blood remains challenging due to their phenotypic heterogeneity and rarity. Label-free microfluidic technologies offer a promising alternative to affinity-based approaches by exploiting intrinsic biophysical differences between cell types. Here, we developed a microfluidic platform for label-free cell separation based on insulator-based dielectrophoresis (iDEP). The microfluidic device employs an array of triangular insulating structures that generate strong electric field gradients in response to an externally applied alternating current (AC) electric field, enabling selective isolation of breast cancer cells from blood cells based on their dielectric properties. Hydrodynamic focusing is used to confine the sample stream and precisely control cell trajectories within the separation region. Numerical simulations were performed to optimize the electric field distribution and fluid flow characteristics within the device. Experimental validation using breast cancer cell lines (mesenchymal-like MDA-MB-231 cells and epithelial-like MCF-7 cells) spiked into peripheral blood mononuclear cells (PBMCs) demonstrates selective dielectrophoretic deflection of cancer cells while PBMCs largely follow the central streamline. The platform achieves recovery rates exceeding 98% and a separation purity above 65% within the optimized operating conditions. The proposed system provides a simple label-free approach to separate heterogeneous cell populations and represents a promising tool for microfluidic liquid biopsy enrichment applications.

## Introduction

Liquid biopsy has emerged as a minimally invasive approach for detection and characterization of cancer-associated biomarkers, including circulating tumor cells (CTCs) ^1,2^. CTCs provide unique insight into tumor heterogeneity, disease progression, and response to treatment ^3,4^. However, the biological and biophysical heterogeneity of CTCs poses a major challenge to their reliable isolation and downstream analysis ^5^. Variations in cellular phenotype, morphology, deformability, and dielectric properties, together with dynamic epithelial–mesenchymal transition (EMT) states, have been frequently associated with metastatic progression, therapeutic resistance, and patient-specific treatment response ^6,7^. Consequently, the isolation and characterization of diverse CTC subpopulations is considered increasingly important for cancer diagnosis and precision oncology applications.

Microfluidic platforms have been extensively explored for the isolation and characterization of CTCs ^8^, owing to their potential for scalable processing, compatibility with clinical workflows, and ability to handle clinically relevant sample volumes. Microfluidic strategies for CTC isolation can broadly be classified into affinity based and label-free approaches ^9^. Affinity-based methods rely on antibody recognition of surface biomarkers, such as epithelial cell adhesion molecule (EpCAM), leukocyte common antigen (CD45), enabling specific and non-specific enrichment of epithelial-like tumor cells ^10^. Representative platforms include the clinically validated CellSearch System ^11^, as well as the AdnaTest ^12^ and PowerMag System ^13^, which utilize immunomagnetic separation principles for CTC enrichment. Despite their clinical utility, these approaches remain inherently biased toward specific phenotypic subpopulations and may fail to capture the full CTC spectrum, such as clinically relevant CTC subpopulations that downregulate epithelial markers during epithelial–mesenchymal transition (EMT) ^14–16^. The importance of addressing CTC heterogeneity has been increasingly emphasized in numerous studies in cancer biology and microfluidics-based diagnostics ^17–22^. In the context of microfluidic technologies, acknowledging and incorporating CTC heterogeneity into device design is therefore essential for achieving improved recovery including clinically potentially relevant sub populations.

Label-free microfluidic platforms enable selective manipulation without molecular tagging; thus, they have the opportunity to become powerful tools for enrichment and isolation of heterogeneous CTCs ^23^ and CTC clusters ^24^. They exploit intrinsic biophysical and electrical differences between cell populations, including size, deformability, density, and dielectric properties ^25^. Among these label-free approaches, dielectrophoresis (DEP) is particularly attractive due to its sensitivity to subtle variations in cellular electrical properties and its capacity for continuous, tunable, and marker-independent cell separation ^26–28^. It can be implemented using either electrode-based or insulator-based architectures. In electrode-based DEP (eDEP) systems, microfabricated metallic electrodes are integrated within the fluidic environment to generate spatially non-uniform electric fields that enable controlled positive or negative DEP responses through adjustment of signal frequency and amplitude ^29–32^. Although this configuration provides precise field modulation, its practical implementation is often constrained by electrode fouling, electrolysis, and long-term material degradation ^33^. These effects can compromise cell viability, limit operational stability, and reduce reproducibility, particularly when processing biologically relevant high conductivity samples such as blood. In contrast, insulator-based DEP (iDEP) uses passive insulating structures within a microchannel to create non-uniform electric fields when a voltage is applied across external electrodes at the ends of the channel ^34–36^. By eliminating the need for patterned electrodes within the fluidic domain, this architecture facilitates the development of robust and cost-effective devices through the use of alternative fabrication methods, including paper-based substrates ^37^ and 3D printing technologies ^38^.

Several microfluidic DEP platforms have previously demonstrated the manipulation and separation of cancer cells from blood cell populations using differences in their dielectric properties ^39–42^. These studies have established the feasibility of label-free enrichment using both electrode-based and insulator-based DEP architectures and have reported promising recovery efficiencies for individual cancer cell lines under controlled experimental conditions. Although phenotypic and biophysical heterogeneity is a defining characteristic of CTC populations in clinical samples, relatively little attention has been paid to designing and evaluating DEP-based microfluidic platforms that are capable of robust, high-recovery enrichment across diverse cancer cell phenotypes. Furthermore, relatively few eDEP ^32,43^ and iDEP ^44^ studies have combined hydrodynamic focusing with geometry-guided design to improve manipulation consistency under continuous-flow conditions. These limitations highlight the need for the development of microfluidic platforms capable of robust, marker-free enrichment across diverse cancer cell phenotypes.

In this study, we demonstrate the selective manipulation and enrichment of heterogeneous breast cancer cell populations from peripheral blood mononuclear cells (PBMCs) using a continuous-flow microfluidic platform based on insulator-based dielectrophoresis. This capability is enabled by the integration of hydrodynamic focusing, which is a versatile tool used in flow cytometry ^45–47^ to align cells within a narrow electric-field interaction zone. Hence, it enhances the consistency of DEP-driven manipulation. To capture clinically relevant phenotypic diversity, we investigate two widely used breast cancer cell models representing mesenchymal-like (MDA-MB-231) ^48,49^ and epithelial-like (MCF-7) ^50,51^ characteristics. The device incorporates ratchet-like insulating structures designed to generate strong localized electric field gradients, enabling modulation of cancer cell trajectories while minimally affecting blood cell populations. Through combined numerical analysis and experimental validation, we show that the platform achieves high-recovery enrichment of cancer cells within heterogeneous suspensions.

## Materials and methods

### Device design and operating principle incorporating DEP theory

The microfluidic device was designed to enable continuous, label-free separation of cancer cells from peripheral blood mononuclear cells (PBMCs) using insulator-based dielectrophoresis (iDEP). As shown in Figures 1a and 1b, the device consists of two inlets (a sample inlet and a sheath flow inlet) and two outlets (a waste outlet and a CTC outlet). A mixture of cells is introduced through the sample inlet. The main microchannel comprises three functional regions: a hydrodynamic focusing section to confine the sample stream, a constriction region incorporating triangular ratchet-like insulating structures to generate non-uniform electric field, and an outlet section for separated stream collection. A sheath flow from side inlet channels is applied at equal rates to narrow the sample stream towards the center of the channel. Hydrodynamic focusing is crucial for maintaining a narrow sample stream inside the constriction channel, ensuring uniform exposure to DEP and keeping PBMCs on the center streamline which routes to the waste outlet (Figure 1c).

**Fig. 1.**
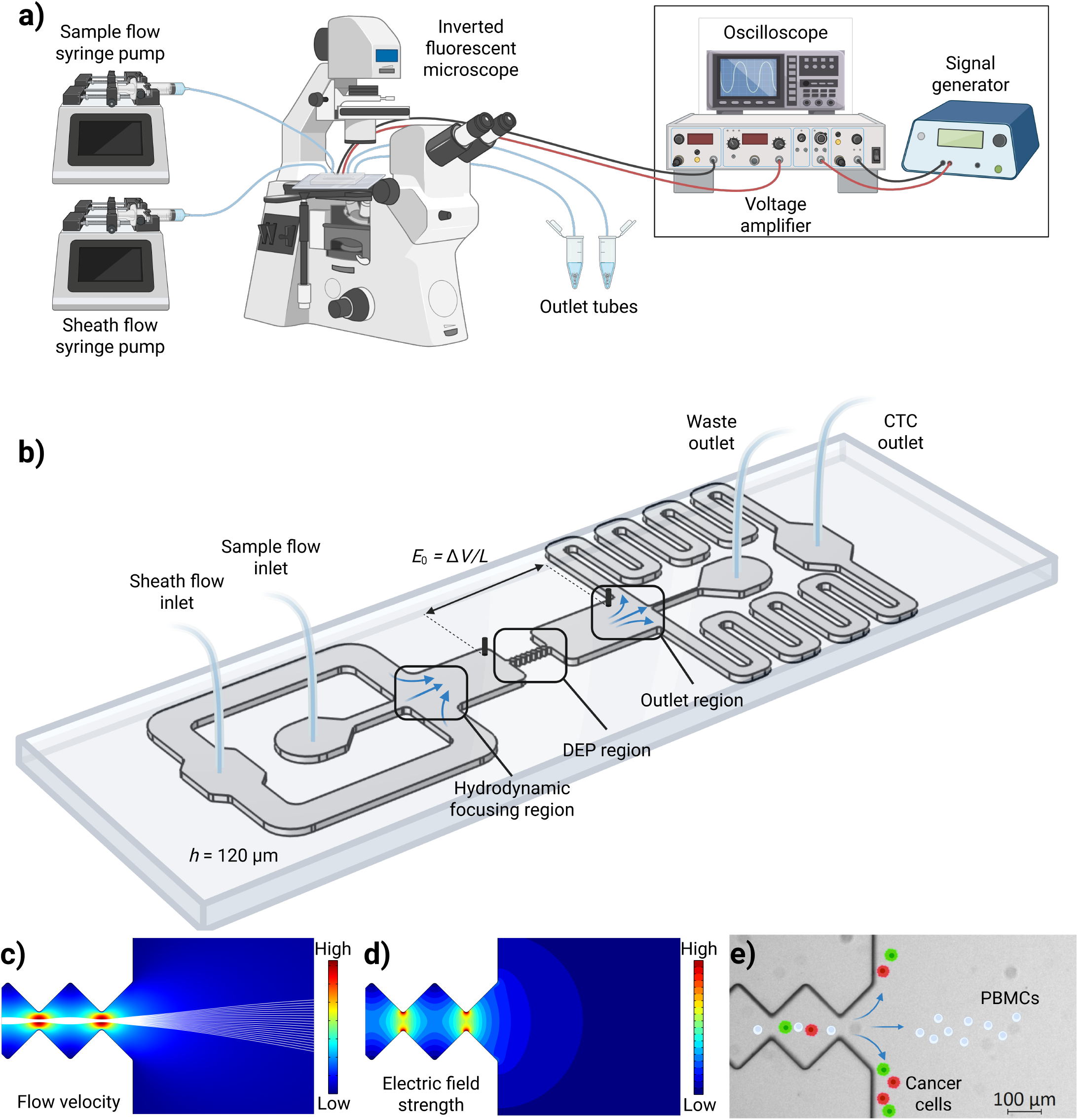
Schematic representation of the microfluidic device and experimental setup, together with the operating principle of iDEP. (a) Fluidic connections are established polytetrafluoroethylene (PTFE) tubing (0.5 mm i.d.; blue), and flow is controlled using two syringe pumps dedicated to sample and sheath streams. The microfluidic chip is mounted on an inverted fluorescence microscope for image acquisition. The electrical setup comprises a signal generator, a voltage amplifier, and an oscilloscope for signal monitoring. (b) The microfluidic device has a channel height of 120 *µ*m, a width of 2.25 mm in the main channel, and a constriction width of 50 *µ*m at DEP region. An electric field, *E*_0_ = Δ*V/L*, is generated across the obstacle array by applying a potential difference, Δ*V* using two platinum electrodes (black cylinders) spaced 5.5 mm apart. (c) Simulated fluid velocity distribution in the constriction channel exit. White streamlines indicate the focused sample stream. The maximum fluid velocity occurs between the tips of the triangular obstacles in the constriction region. (d) Simulated electric field distribution illustrating the high-field regions formed at the tips of the insulating triangular structures due to the non-uniformity of the applied electric field. (e) Schematic illustration of particle manipulation in the actual PDMS channel. The dielectrophoretic force acts perpendicular to the flow direction; cancer cells (green and red) experiencing pDEP are attracted toward the high-field regions at the obstacle tips and are deflected toward trajectories closer to the sidewalls. At the selected operating frequency, PBMCs (white) experience limited DEP forces and remain along the central streamline. In the absence of DEP forces, all cells follow the hydrodynamically focused center path and exit through the waste outlet (not shown here). Cell trajectories are determined by the balance between hydrodynamic drag and dielectrophoretic forces, which can be tuned through the applied voltage and flow rates.

As the focused flow enters the constriction channel, the constrictions generate strong electric field gradients when an AC voltage is applied across the channel (Figure 1d). Platinum electrodes (0.38 mm diameter; Goodfellow, PT00-WR-000143) are inserted through designated through-holes on both sides of the constriction region, spaced 5.5 mm apart, allowing voltage application without embedding metal micro-electrodes within the actual DEP manipulation region. The non-uniform electric field generates DEP forces that redirect positively DEP-experienced cancer cells towards the tips. As a result, they are moved onto streamlines directed toward the cancer cell outlet. In contrast, PBMCs experience weaker, and negligible DEP forces, remain in their original streamlines and proceed along the central axis, directed towards the waste outlet (Figure 1e). This allows sorting into distinct outlets based on cells’ DEP response.

The time-averaged DEP force *F*_DEP_, acting on a spherical particle of radius *r*, suspended in a medium with a relative permittivity *ε*_m_, is given by:

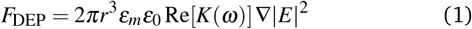

where *ε*_0_ is permittivity of vacuum, *E* is the amplitude (rms) of the electric field, ∇ is the gradient operator, *ω* = 2*π f* is the angular frequency of the applied signal, *f* is the frequency of the applied field, and Re[*K*(*ω*)] is the real part of the Clausius–Mossotti factor. This factor is defined as

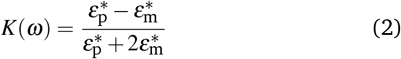

with the complex permittivities 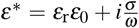 of the particle (subscript p) and the surrounding medium (subscript m). Here, *ε*_r_ and *σ* are the relative permittivity and conductivity of the particle and medium, respectively, *ε*_0_ is the permittivity of free space, and 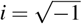 is the imaginary number. The sign and magnitude of Re[*K*(*ω*)] determine the direction and strength of the DEP force. When Re[*K*(*ω*)] *>* 0, positive DEP (pDEP) occurs, cells are attracted to regions of high electric field intensity. When Re[*K*(*ω*)] *<* 0, negative DEP (nDEP) occurs, cells are repelled from high-field regions and migrate to low electric field regions. At Re[*K*(*ω*)] = 0, zero DEP occurs, the particle does not experience a net DEP force, although it is still subject to fluidic drag.

Due to the frequency dependence of the complex permittivity, the Clausius–Mossotti factor and therefore the DEP behavior of a cell (pDEP or nDEP) depend on the frequency of the applied field. While homogeneous particles usually experience only a single change in the DEP behavior with frequency (one *dispersion*), mammalian cells are usually modeled using the single-shell model and experience two dispersions, one in the kHz range and one in the MHz range ^26,52–54^. The frequency of the first dispersion is slightly lower for most cancer cells than for white blood cells, as it is inversely related to the cell membrane capacitance ^55^. Cancer cells typically exhibit higher membrane capacitance due to their more complex and highly folded (wrinkled) membrane morphology, resulting in a lower first dispersion frequency ^55,56^.

Figure 2 shows the value of Re[*K*(*ω*)] for white blood cells and two breast cancer cell lines (MCF-7 and MDA-MB-231) at a medium conductivity of 2 mS/m. Low-conductivity media are commonly employed in DEP systems to reduce Joule heating associated with the application of high electric fields ^57^. 8 kHz was selected as the operating frequency for selective pDEP-based isolation of cancer cells from PBMC populations. This frequency is consistent with the conductivity dependent crossover behavior of white blood cells reported in a previous study ^58^, where a crossover frequency of 271 kHz was observed at a medium conductivity of 70 mS/m, corresponding to approximately 8 kHz under the lower conductivity conditions (2 mS/m) used in the present study. Under these conditions, cancer cells exhibit pDEP behavior, whereas most white blood cells display only weak pDEP (with Re[*K*(*ω*)] values three-fold lower than those of the cancer cells) or transition toward nDEP.

**Fig. 2.**
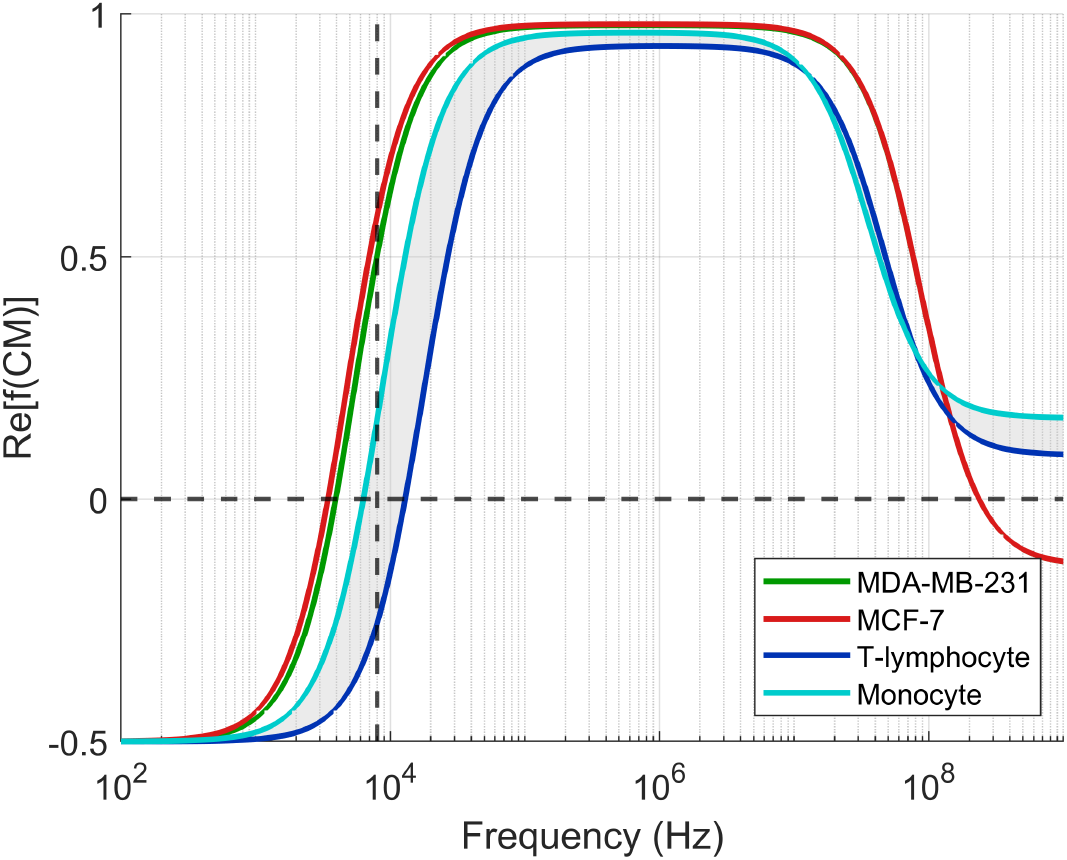
Real part of the Clausius–Mossotti spectra of the MDA-MB-231, MCF-7 breast cancer cell lines ^59^, and major PBMC subpopulations ^60^ calculated at a medium conductivity of 2 mS/m using MyDEP ^61^. The grey shaded region indicates the DEP response envelope between T-lymphocytes and monocytes, the dominant PBMC populations. The dashed vertical line marks the selected operating frequency (8 kHz), while the dashed horizontal line corresponds to Re[*K*(*ω*)] = 0. At 8 kHz, cancer cells exhibit pDEP having Re[*K*(*ω*)] *>* 0.5, whereas PBMC subpopulations show weak pDEP (Re[*K*(*ω*)] *≈* 0.16) or nDEP (Re[*K*(*ω*)] *≈ −*0.25).

At the end of the main channel, the outlets are divided into a central waste outlet, which collects the unmanipulated PBMCs, and two side outlets that merge to collect the deflected cancer cells. The effectiveness of this mechanism relies on a careful balance between fluidic drag and DEP force. Hydrodynamic focusing plays a crucial role in confining the sample flow, ensuring that PBMCs remain within the central streamline and reach the waste outlet, while manipulated cancer cells are displaced toward the sidewalls and sorted accordingly.

To verify that the proposed device geometry and operating conditions are sufficient for effective dielectrophoretic manipulation, two-dimensional and three-dimensional finite-element simulations were performed. Simulations were used to evaluate the flow velocity distribution, electric field strength, electric field gradient, and the uniformity of these fields throughout the channel height. Details of the computational model, boundary conditions, mesh, and representative simulation results are provided in the Supplementary Information S1 (Figures S1 and S2).

### Microfluidic device fabrication and experimental setup

The microfluidic devices were fabricated using standard soft lithography techniques ^62^. The master mold was fabricated on a 4 inch silicon wafer in a cleanroom environment using a *µ*MLA Laser Writer (Heidelberg Instruments). The resulting microfluidic channel height was 120 ± 5 *µ*m. Polydimethylsiloxane (PDMS, Sylgard 184, Dow Corning) was mixed in a 10:1 ratio (base to curing agent), degassed under vacuum, and poured over the mold. After curing at 70°C for 2 hours, the PDMS layer was peeled off and access holes for inlets, outlets, and electrode connections were punched using biopsy punches. The PDMS device was irreversibly bonded to a glass slide following oxygen plasma treatment using a plasma cleaner (PDC-002-CE, Harrick Plasma, USA) to seal the microchannels. The completed chip was placed in an oven at 70°C for at least 30 minutes to enhance bonding strength before use.

The experimental setup (Figure 1a) consisted of two syringe pumps (Pump 11 Elite, Harvard Apparatus, USA) that independently drove the sample and sheath flows into their designated inlets at controlled flow rates. The sample and sheath flow rates were set to 5 *µ*L min^*−*1^ and 10 *µ*L min^*−*1^, respectively, corresponding to a sheath-to-sample flow rate ratio of 2:1. This flow configuration was selected to achieve stable hydrodynamic focusing of the sample stream before it enters the DEP region. An AC signal was applied across the device electrodes using a signal generator (Rigol DG4062, Rigol Technologies, PRC), with the operating frequency and voltage selected based on the simulations described in the Supplementary Information. The signal was subsequently amplified 200-fold using a high-voltage amplifier (Trek PZD700A, Advanced Energy Industries, USA). The generated and amplified signals were monitored using an oscilloscope (Rigol DS7014, Rigol Technologies, PRC) to verify waveform integrity and voltage amplitude.

### Microscopy setup, video recording and post processing

Real-time monitoring of cell motion was performed using an inverted fluorescence microscope (Axio Observer Z1/7, Zeiss GmbH, Germany) equipped with a Colibri 7 illumination system. For fluorescence imaging of GFP, mCherry, and Calcein Blue, LED modules with peak wavelengths of 475 nm, 567 nm, and 385 nm were used in combination with 38 HE GFP (ex: 488 nm, emm: 509 nm), 56 HE GFP/DsRed (ex: 587 nm, emm: 610 nm), and 49 DAPI (ex: 400 nm, emm: 452 nm) filter sets, respectively. Time-lapse image sequences were acquired using a 10× air objective (Plan-Apochromat 10x/0.45 M27) and an ORCA Flash 4.0 V2 sCMOS camera (Hamamatsu Photonics, Japan) at a resolution of 2048 × 2048 pixels.

For each experimental condition, 50 fluorescence frames were recorded with an exposure time of 5 ms using Zen 3.13 software and stored in .czi format. The acquired images were subsequently processed in ImageJ (v1.54p, National Institutes of Health, USA). A detailed image processing workflow is provided in the Supplementary Information S2. The results of the post processing and cell counting are presented in the results and discussion section.

### Cell culture, PBMC isolation, and sample preparation

Two malignant human breast cancer cell lines were used in the experiments: MDA-MB-231 cells expressing LifeAct-GFP and MCF-7 cells constitutively expressing cytoplasmic mCherry. As these cell lines were genetically modified to express fluorescent markers, no additional staining was required. They were cultured in Dulbecco’s Modified Eagle Medium (DMEM, Gibco) supplemented with 5 % fetal bovine serum (FBS, Gibco). Cells were maintained at 37 °C in a humidified incubator with 5 % CO_2_ and subcultured at least twice per week. All experiments were conducted using cells with passage numbers not exceeding 20 (P20) to minimize passage-related changes in cellular characteristics. Prior to each experiment, the average cell diameter of both cell lines was measured to be approximately 14 *µ*m using an automated cell counter (TC20, Bio-Rad Laboratories, Inc., USA).

PBMCs were isolated from whole blood collected from healthy human donors at Erasmus MC, Rotterdam, following ethical approval and informed consent. Whole blood was diluted 1:1 with sterile phosphate-buffered saline (PBS) and gently layered over Ficoll-Paque Plus (GE Healthcare) in 50 mL conical tubes. Samples were centrifuged at 400 × g for 30 minutes at room temperature with the brake off. After centrifugation, the buffy coat containing the PBMC layer was carefully collected, washed twice with PBS, and centrifuged at 300 × g for 10 minutes to remove residual Ficoll and plasma components. The PBMC were then stained with either Calcein Green or Calcein Blue (Thermo Fisher Scientific, Waltham, MA) at a final concentration of 2 µM to enable fluorescent labeling depending on the experiment type. Following staining, the cells were washed with PBS, and the resulting cell pellet was resuspended in DEP buffer for subsequent microfluidic experiments. Cell concentration and viability were assessed using trypan blue exclusion and an automated cell counter (TC20, Bio-Rad Laboratories, Inc., USA).

To ensure appropriate dielectric contrast and maintain cell viability, a low-conductivity DEP buffer was prepared by dissolving 8.5 % (w/v) sucrose and 0.3 % (w/v) dextrose in deionized (DI) water. A small volume of 1× phosphate-buffered saline (PBS) was added to adjust the conductivity to the target value of 2 mS/m, which is suitable for effective DEP operation. The buffer was sterile-filtered using a 0.22 *µ*m filter and was used fresh for each experiment.

Prior to each experiment, cancer cells and PBMCs were washed twice with DEP buffer and resuspended at the desired concentrations. For single-cell-type experiments, MDA-MB-231 and MCF-7 cells were prepared separately at a concentration of 1 *×* 10^6^ cells mL^*−*1^ in DEP buffer. For co-culture experiments, MDA-MB-231 and MCF-7 cells were mixed at a 1:1 ratio while maintaining a total cancer cell concentration of 1 *×* 10^6^ cells mL^*−*1^. To evaluate separation performance under biologically relevant conditions, spike-in PBMC samples were prepared by mixing cancer cells and PBMCs at a 1:1 ratio, corresponding to final concentrations of 1 *×* 10^6^ cancer cells mL^*−*1^ and 1 *×* 10^6^ PBMCs mL^*−*1^. For all experimental conditions, 1 mL of sample suspension was prepared and introduced into the microfluidic device.

## Results and discussion

### Trajectory modulation of heterogeneous cancer cells under iDEP actuation

To evaluate the separation capability of the iDEP platform, the trajectory behaviour of cells was first investigated under conditions with and without an applied electric field in the DEP region. Hydrodynamic focusing confined MDA-MB-231 cells, MCF-7 cells, and PBMCs to the central streamline of the main microchannel prior to entering the DEP region consisting of a constriction channel incorporating triangular insulating sidewall structures. In the absence of an electric field, all cell populations predominantly followed trajectories toward the waste outlet after exiting the DEP region. This behavior indicates that hydrodynamic drag forces govern cell motion when electric field gradients are absent (Fig. 3a–d).

**Fig. 3.**
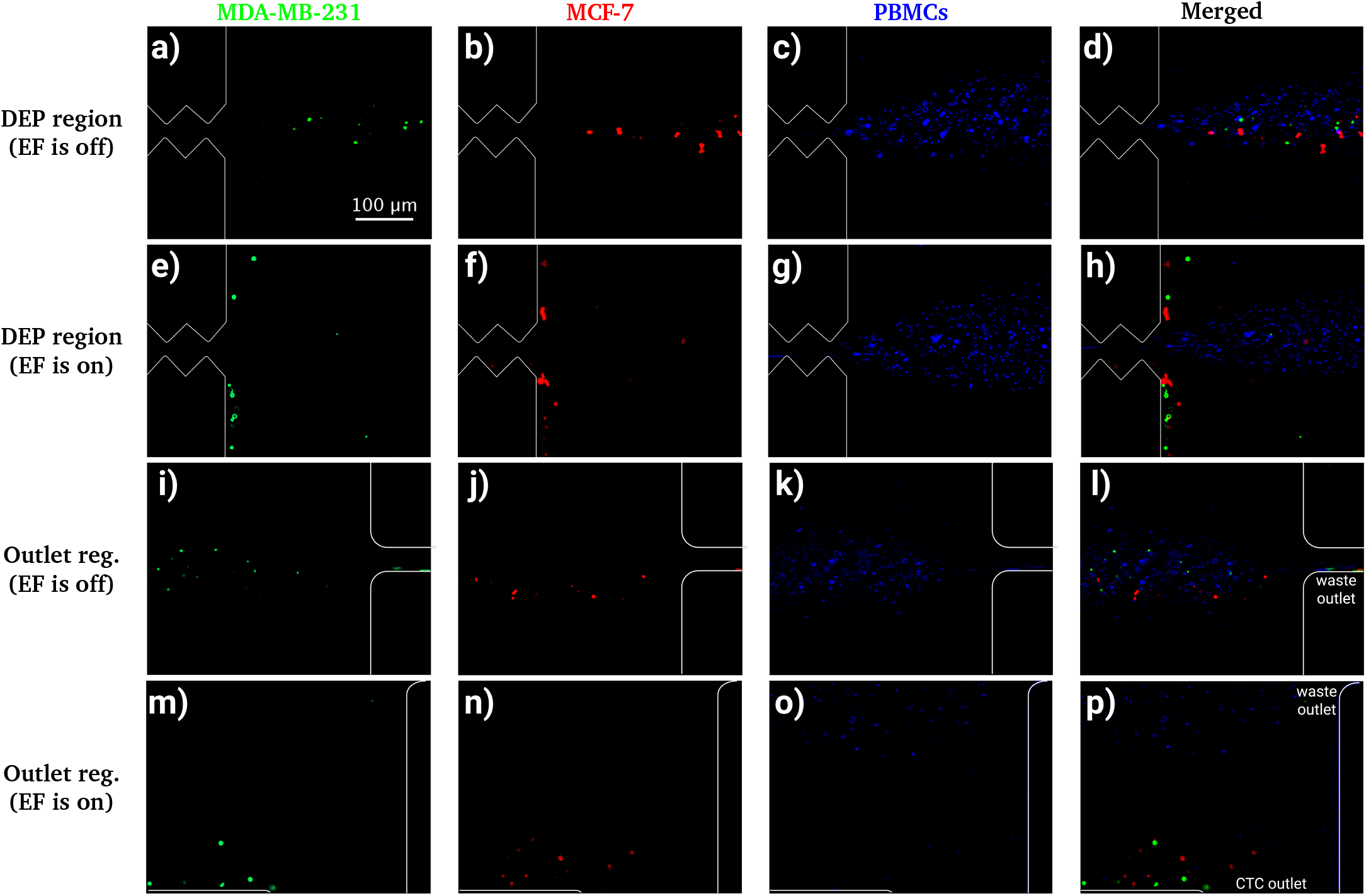
Trajectory modulation of heterogeneous cancer cells and PBMCs under hydrodynamic focusing and dielectrophoretic (DEP) actuation for solo experiments. Following hydrodynamic focusing, cells are confined to the central streamline upon exiting the constriction region and predominantly follow trajectories toward the downstream waste outlet in the absence of an electric field (EF off). Fluorescence microscopy images showing the displacement behavior of (a) MDA-MB-231-LifeAct GFP breast cancer cells, (b) MCF-7-mCherry breast cancer cells, (c) PBMCs stained with Calcein Green, and (d) merged fluorescence images under EF-off conditions. Under these conditions, all cell populations remain broadly distributed around the central streamline and are mainly directed toward the waste outlet. Upon application of an AC electric field (210 *V*_RMS_, 8 kHz), cancer cells experience positive dielectrophoresis (pDEP) near the insulating obstacle tips, resulting in lateral migration. Fluorescence images of (e) MDA-MB-231 and (f) MCF-7 cells demonstrate DEP-induced displacement toward streamlines closer to the channel sidewalls, ultimately redirecting the cells into the designated CTC outlet streams. (g) PBMC trajectory behavior under identical DEP conditions shows only weak lateral displacement near the constriction exit, indicating a limited dielectrophoretic response. (h) Merged fluorescence images under DEP actuation highlight the distinct dielectrophoretic responses between cancer cells and PBMCs. Fluorescence images acquired at the outlet region further demonstrate the separation outcome. Under EF-off conditions, (i) MDA-MB-231 cells, (j) MCF-7 cells, and (k) PBMCs predominantly follow the waste outlet route, as shown in the merged image in (l). When the electric field is activated, (m) MDA-MB-231 and (n) MCF-7 cells are redirected toward the CTC outlet streams (only the bottom CTC outlet is shown). In contrast, PBMCs largely remain in the waste outlet stream, although a fraction of the population follows trajectories closer to the sidewalls and enters the CTC outlet streams, contributing to reduced purity in the collected cancer cell fractions. The corresponding merged fluorescence image is shown in (p). All fluorescence images were acquired at the same magnification. Fluorescence signals were recorded in grayscale and are displayed using consistent pseudocolors assigned during image processing (green for MDA-MB231 cells, red for MCF7 cells, and blue for PBMCs) to facilitate visualization and comparison of the different cell populations. Flow direction is from left to right, and the scale bar in panel (a) applies to all panels. While Figure 3 illustrates representative snapshots from the solo experiments, the dynamic behavior observed during co-culture and PBMC spike-in experiments is presented in Supplementary Videos S1–S8.

Cancer cells experienced positive dielectrophoretic forces near the tips of the insulating ratchet structures when an AC electric field (210 V_RMS_, 8 kHz) was applied. As a result, the manipulated cells were laterally displaced toward streamlines closer to the channel sidewalls while traversing the DEP region (Fig. 3e & 3f). In contrast, PBMCs exhibited negligible lateral displacement and largely maintained their original streamlines (Figure 3g). This enables effective spatial separation between cancer and blood cell populations. Comparable trajectory modulation was observed for both epithelial-like (MCF-7) and mesenchymal-like (MDA-MB-231) phenotypes, demonstrating robust DEP-driven manipulation across heterogeneous cancer cell types.

At the outlet region, fluorescence imaging further confirmed the separation behavior observed in the DEP region. In the absence of an applied electric field, MDA-MB-231 cells, MCF-7 cells, and PBMCs predominantly followed the hydrodynamically focused central streamline and exited through the waste outlet (Fig. 3i–l). This observation is consistent with the dominance of hydrodynamic drag forces under EF-off conditions, where no lateral trajectory modulation occurs.

Upon application of the AC electric field, cancer cells underwent lateral displacement due to positive dielectrophoretic forces generated near the insulating ratchet structures. Consequently, both MDA-MB-231 and MCF-7 cells were redirected toward streamlines near the channel sidewalls and subsequently collected at the designated CTC outlet (Fig. 3m & 3n). In contrast, the majority of PBMCs remained within the central streamline and continued toward the waste outlet, although a fraction of the population exhibited lateral displacement and entered the CTC outlet streams (Fig. 3o). This partial co-enrichment of PBMCs is attributed to overlapping dielectric and biophysical characteristics between certain PBMC subpopulations and the target cancer cells, contributing to reduced purity in the collected fractions. The merged fluorescence image in Figure 3p further demonstrates the selective isolation of heterogeneous cancer cell populations from PBMCs. Representative real-time recordings of the co-culture and spike-in PBMC experiments under both EF-on and EF-off conditions are provided in Supplementary Videos S1– S4 and S5–S8, respectively.

One potential limitation of DEP-based systems is Joule heating arising from electric current passing through the conductive medium, which may adversely affect cell viability at high electric field strengths and solution conductivities. In the present study, experiments were performed using a low-conductivity DEP buffer (2 mS m^*−*1^), thereby minimizing Joule heating. Throughout the experiments, no evidence of bubble formation, flow instability, or obvious loss of cellular fluorescence was observed following passage through the DEP region, suggesting that thermal effects were not sufficiently severe to compromise the performed experiments.

### Quantitative analysis of recovery and purity

To quantify cell distributions and sample purity, outlet regions were considered. Recorded image frames were imported into ImageJ, where a semi-automated fluorescent cell counting workflow was applied to quantify particles within identical regions of interest (ROIs) positioned downstream of the bifurcation. This workflow consisted of standardized thresholding, ROI selection, and size-based particle detection steps applied identically across all frames, as summarized in Figure 4 and the Supplementary Information. Particle counts obtained for each frame and outlet were averaged to determine the mean cell occupancy per frame.

**Fig. 4.**
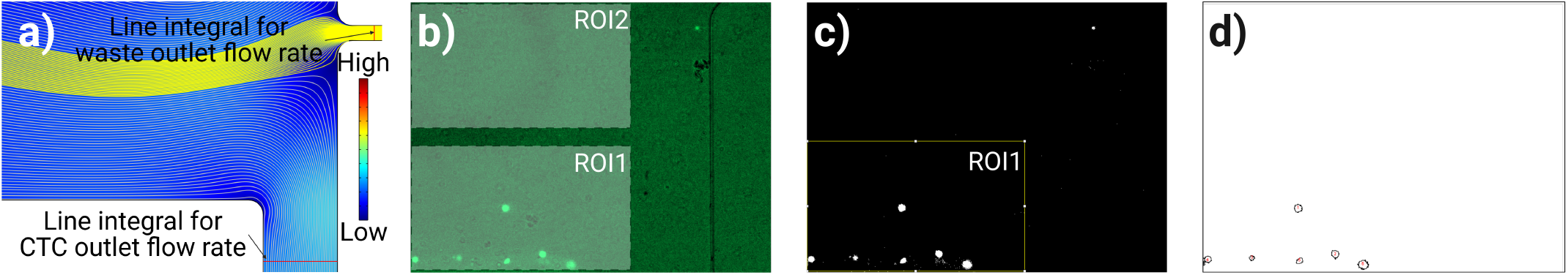
Cell counting methodology. (a) COMSOL Multiphysics 6.2 (COMSOL AB, Sweden) flow simulations showing the separation of sample and waste streams at the outlet region. Waste fluid is directed toward the waste outlet (yellow streamlines), whereas sample fluid exits through the upper and bottom CTC outlets (gray streamlines). Outlet flow rates were obtained by integrating the velocity field along vertical cut lines (line integrals) at each outlet. These values were subsequently used to calculate cell fluxes based on the average cell occupancies per frame (*n*_1,avg_ and *n*_2,avg_) measured within ROI1 and ROI2 in Fig.4b, separately. (b) Representative fluorescence image of the same outlet region used for experimental quantification of total MDA-MB-231 cell occupancies (*N*_1_ and *N*_2_) within ROI1 and ROI2 over 50 consecutive frames. (c) Binary threshold image generated in ImageJ showing the defined ROIs used for automated cell detection. Object selection criteria included minimum diameter and circularity thresholds. (d) Final cell detection result, where detected cells are indicated by circular outlines.

Because instantaneous ROI occupancies depend on local residence time, raw occupancies could not be interpreted directly as particle fractions due to unequal outlet flow conditions. In the present microfluidic system, varying sheath and sample flow rates resulted in non-uniform outlet flow distributions, thereby affecting cell residence times within the monitored ROIs. Hence, occupancies were converted into outlet-specific cell fluxes. Local velocities and volumetric flow rates at the ROIs were obtained from laminar-flow COMSOL simulations using the experimental geometry and boundary conditions (Figure 4a). Outlet flow rates were determined by integrating the velocity field along vertical cross-sections (line integrals) at each outlet. Detailed outlet flow-rate calculations and corresponding COMSOL expressions are provided in Table S1 in the Supplementary Information. Cell fluxes were subsequently calculated by combining the average number of detected cells per frame within each ROI with the corresponding outlet flow rate.

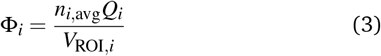

where Φ_*i*_ is the cell flux at outlet *i* (cells/min), *n*_*i*,avg_ is the average number of detected cells per frame within ROI_*i*_, *Q*_*i*_ is the corresponding outlet volumetric flow rate (*µ*l/min), and *V*_ROI,*i*_ is the effective observation volume (*µ*l) of the ROI.

Recovery efficiency was calculated by combining the fluxes from the two CTC outlet streams and normalizing them to the total target-cell flux collected from all outlets:

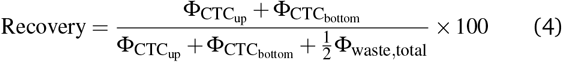

Since the waste outlet was visible in both outlet-region recordings, cells entering it were counted twice during image analysis. To avoid overestimating the waste fraction, the total measured waste flux was divided by two before calculating recovery efficiency. Recovery was therefore calculated as the combined cancer-cell flux collected from the two CTC outlets relative to the corrected total cancer-cell flux across all outlet streams.

For spike-in PBMC experiments, purity was calculated as the fraction of cancer-cell flux within the combined CTC outlet streams relative to the total collected cell flux in these outlets:

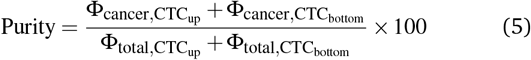

Experimentally, fluorescence image sequences of the outlet region were acquired and used to quantify total cell counts (*N*_1_ and *N*_2_) over 50 consecutive frames. Image analysis was performed in ImageJ, where binary thresholding was applied to generate segmentation masks for automated cell detection (Figure 4c). Regions of interest corresponding to each outlet stream were defined, and object selection was performed according to the applied analysis protocol (described in the Supplementary Information), using size (minimum and maximum area) and circularity criteria to exclude debris and noise. The final detection output consisted of identified cells within each ROI separately (Figure 4d), enabling consistent and reproducible quantification of outlet-specific cell distributions.

Recovery efficiency as sample complexity increases is shown in Figure 5. First, single-cell-type suspensions of MDA-MB-231 and MCF-7 were used to evaluate the performance under simplified conditions. High and consistent recovery rates were obtained for both cell lines, with values of (98.3 *±* 0.75)% and (99.5 *±* 0.50)%, respectively. Then, to assess the impact of inter-cell-type interac-

**Fig. 5.**
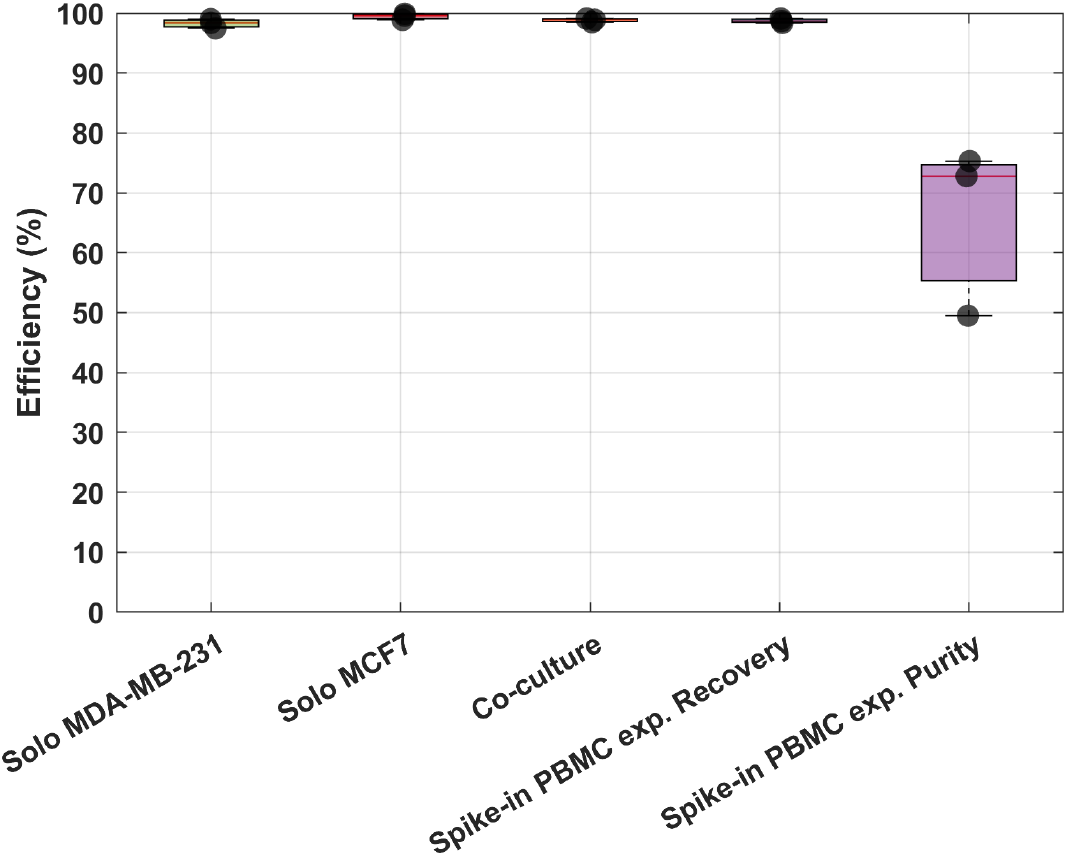
Recovery performance across increasing sample complexity and corresponding purity in spike-in PBMC conditions. Box plots summarizing the recovery efficiency of MDA-MB-231 and MCF-7 under three experimental conditions of increasing biological complexity. Solo experiments represent single-cell-type suspensions of each cell line processed independently. Co-culture experiments correspond to a 1:1 mixture of MDA-MB-231 and MCF-7 cells, enabling assessment of recovery performance in the presence of inter-cell-type interactions. Spike-in PBMC experiments represent a complex biological matrix in which cancer cells are spiked into PBMCs, mimicking clinically relevant conditions. For the spike-in PBMC case, both recovery (fraction of target cancer cells collected at the designated outlet relative to the initial input) and purity (fraction of recovered cells identified as target cancer cells relative to total collected cells) are reported.

tions, experiments were subsequently performed using a 1:1 co-culture of the two cell lines. Under these conditions, the recovery remained comparably high ((98.8 *±* 0.31)%), indicating that the presence of a second cancer cell population did not adversely affect manipulation efficiency. Further evaluation was performed by spiking cancer cells into PBMC suspensions at a 1:1 cancer cell-to-PBMC ratio, thereby introducing a heterogeneous cellular background and enabling assessment of the platform under increased sample complexity. The cancer cell recovery rates in the samples with cancer cells spiked into PBMCs ((98.7 *±* 0.35)%) were consistent with both the solo and co-culture experiments. No statistically significant differences were observed in recovery rates among solo, co-culture, and spike-in PBMC conditions (Welch’s t-test, p > 0.05), indicating that separation performance is maintained as sample complexity increases. The comparable recovery efficiencies obtained for both epithelial-like (MCF-7) and mesenchymal-like (MDA-MB-231) phenotypes also demonstrate the platform’s robustness across heterogeneous cancer cell populations.

In addition to recovery, the purity of collected samples was evaluated in spike-in PBMC experiments to assess separation selectivity under clinically relevant conditions. While recovery remained consistently high ((98.7 *±* 0.35)%), purity was substantially lower and exhibited greater variability, with a value of ((65.9 *±* 14.1)%). The reduced purity was primarily attributed to the co-enrichment of non-target PBMCs, particularly monocytes, which exhibit electrokinetic characteristics that partially overlap with those of the target cancer cells under the applied operating conditions. Furthermore, variations in monocyte content between biological replicates contributed to the observed variability in purity. The observed trade-off between recovery and purity is a characteristic of CTC isolation approaches ^63^. Preservation of the target cell population is generally prioritized over complete depletion of background cells because target cells are often rare and biologically valuable. Consequently, high recovery is considered more critical than achieving maximal purity, as residual background cells can frequently be addressed during downstream analyses, whereas lost target cells cannot be recovered ^64^.

## Conclusions

In this study, we developed and experimentally validated a microfluidic platform for label-free manipulation and enrichment of heterogeneous breast cancer cell populations using insulator-based dielectrophoresis. The integration of hydrodynamic focusing with a ratchet-like insulating obstacle array enabled the generation of localized electric field gradients that selectively deflected cancer cells while allowing peripheral blood mononuclear cells to predominantly follow the central streamline. Numerical modeling supported device optimization and provided insight into the interplay between hydrodynamic drag and dielectrophoretic forces governing cell trajectories. Under optimized operating conditions, the system achieved high recovery efficiencies and clinically relevant purity levels, demonstrating its capability for continuous and marker-independent cell enrichment. These results highlight the potential of iDEP-based microfluidic architectures as robust and scalable tools for unbiased circulating tumor cell enrichment, with promising implications for next-generation liquid biopsy workflows and precision oncology applications.

## Author contributions

Conceptualization, G.Ö., and P.E.B.; investigation, G.Ö., I.N.U., E.Y., and J.K.; methodology, G.Ö., I.N.U., J.K., D.B., G.R.P., and P.E.B.; validation, G.Ö., I.N.U., and E.Y.; formal analysis, G.Ö. and G.R.P.; writing—original draft preparation, G.Ö.; writing—review and editing, G.Ö., J.K., J.W.M.M., P.tD., G.R.P., and P.E.B; visualization, G.Ö., I.N.U, and E.Y.; resources, D.B., K.D., J.W.M.M, P.tD., and P.E.B.; supervision, G.Ö., K.D., P.tD., G.R.P., and P.E.B.; funding acquisition, J.W.M.M., P.tD., and P.E.B. All authors have read and agreed to the published version of the manuscript.

## Data availability

The data supporting the findings of this study are available in the Supplementary Information (SI). The complete experimental datasets, raw and processed data, MATLAB codes, and supplementary videos have been deposited in the 4TU.ResearchData repository and are openly available at https://doi.org/10.4121/464f2d63-757c-4351-8a99-0eca7c427337.

## Acknowledgements

This work was supported by the Netherlands Organization for Health Research and Development (ZonMw, project no. 09120012010061). Artificial intelligence (AI)-assisted tools (ChatGPT, OpenAI) were used for language refinement and editorial support during manuscript preparation. All scientific content, data analysis, interpretations, and conclusions were reviewed and approved by the authors.

