## Supplementary Information file for "Label-free Isolation of Heterogeneous Breast Cancer Cell Populations via Insulator-Based Dielectrophoresis^†^"

### Supplementary Videos

The supplementary videos provide representative real-time fluorescence microscopy recordings of the co-culture and spike-in PBMC experiments under both EF-on and EF-off conditions. Videos were acquired at the DEP and outlet regions to illustrate the dynamic trajectory modulation and separation of heterogeneous cancer cell populations.

- **Video S1.** MDA-MB-231-LifeAct GFP and MCF-7-mCherry co-culture experiment recorded at the DEP region under EF-on conditions.
- **Video S2.** MDA-MB-231-LifeAct GFP and MCF-7-mCherry co-culture experiment recorded at the DEP region under EF-off conditions.
- **Video S3.** MDA-MB-231-LifeAct GFP and MCF-7-mCherry co-culture experiment recorded at the outlet region under EF-on conditions.
- **Video S4.** MDA-MB-231-LifeAct GFP and MCF-7-mCherry co-culture experiment recorded at the outlet region under EF-off conditions.
- **Video S5.** Spike-in PBMC experiment (MDA-MB-231-LifeAct GFP, MCF-7-mCherry, and PBMCs) recorded at the DEP region under EF-on conditions.
- **Video S6.** Spike-in PBMC experiment (MDA-MB-231-LifeAct GFP, MCF-7-mCherry, and PBMCs) recorded at the DEP region under EF-off conditions.
- **Video S7.** Spike-in PBMC experiment (MDA-MB-231-LifeAct GFP, MCF-7-mCherry, and PBMCs) recorded at the outlet region under EF-on conditions.
- **Video S8.** Spike-in PBMC experiment (MDA-MB-231-LifeAct GFP, MCF-7-mCherry, and PBMCs) recorded at the outlet region under EF-off conditions.

### S1. Numerical simulation and model validation

Numerical simulations were performed to verify the hydrodynamic and electrical operating conditions of the final microfluidic device and to support the interpretation of the experimental results.

Finite-element simulations were performed using COMSOL Multiphysics 6.2 (COMSOL AB, Sweden) to evaluate fluid flow and electric field distributions within the final microfluidic device geometry. The simulations employed two coupled physics interfaces: Creeping Flow and Electric Currents (frequency domain). Material properties and boundary conditions were selected to reproduce the experimental operating conditions.

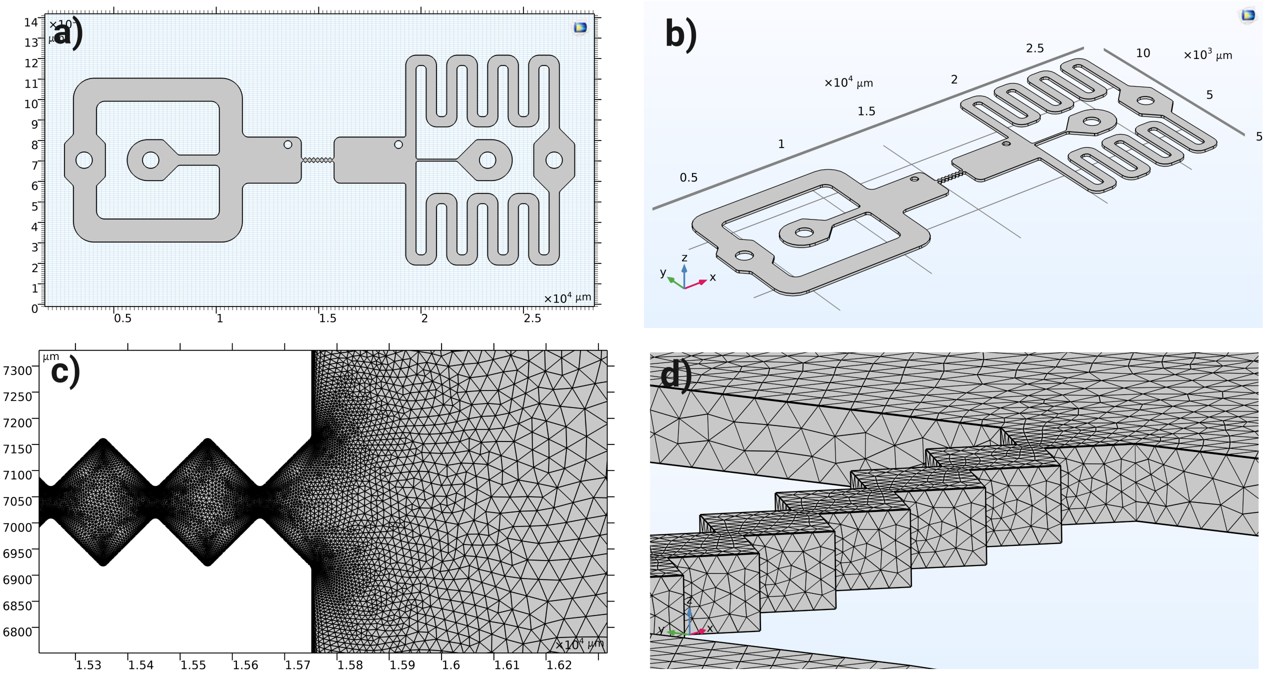


**Fig. S1** Computational models used for finite-element simulations. (a) Two-dimensional axisymmetric geometry of the microfluidic device used for fluid flow, electric field, and particle-tracing simulations. Boundary conditions applied to the inlet, outlet, channel walls, and electrodes are indicated. (b) Three-dimensional geometry used to evaluate the uniformity of the electric field distribution across the 120 $\mu$m channel height within the DEP region. (c) User-controlled computational mesh of the two-dimensional model generated using free triangular elements with the predefined Extra Fine Fluid Dynamics mesh sequence. Three boundary-layer elements with a thickness factor of 2.6 were applied along all channel walls to improve the resolution of near-wall flow gradients. Local mesh refinement around the constriction region and insulating obstacle tips enables accurate resolution of the steep electric field gradients responsible for dielectrophoretic manipulation.

Fluid flow was modeled using the Creeping Flow interface under steady-state, incompressible conditions. The sample and sheath inlet boundary conditions were prescribed as volumetric flow rates of 5 and 10 $\mu$L min$^{-1}$, respectively. Because the computational model was two-dimensional, the prescribed volumetric flow rates were converted to equivalent average inlet velocities by assuming the inlet to represent the lateral surface of a cylindrical channel. Accordingly, the effective inlet area was approximated as $A = 2\pi rh$, where $r = 400$ $\mu$m is the inlet radius and $h = 120$ $\mu$m is the channel height. A pressure boundary condition ($P = 0$ Pa) was applied at the outlet, while no-slip boundary conditions were imposed on all channel walls. The applicability of the Creeping Flow interface was verified by estimating the characteristic Reynolds number of the flow. The Reynolds number was calculated as:

$$\mathrm{Re} = \frac{\rho U D_h}{\mu}, \tag{1}$$

where $\rho$ is the fluid density, $U$ is the average inlet velocity, $D_h$ is the hydraulic diameter of the inlet channel, and $\mu$ is the dynamic viscosity of the DEP buffer. The average inlet velocity was determined from the total volumetric flow rate according to:

$$U = \frac{Q}{A}, \tag{2}$$

where $Q$ is the total inlet flow rate and $A$ is the effective inlet area used to prescribe the inlet boundary condition. The hydraulic diameter of the rectangular main channel was calculated as:

$$D_h = \frac{2wh}{w+h}, \quad (3)$$

where $w = 2250$ $\mu$m is the main channel width and $h = 120$ $\mu$m is the channel height. Using a total flow rate of 15 $\mu$L min$^{-1}$, DEP buffer fluid properties ($\rho = 1000$ kg m$^{-3}$ and $\mu = 1.0 \times 10^{-3}$ Pa s), and the calculated average inlet velocity, the Reynolds number was estimated to be approximately 0.21. This confirms that viscous forces dominate over inertial effects and validates the use of the Creeping Flow interface for the present simulations.

The electric field distribution was calculated using the Electric Currents interface in the frequency domain. One electrode was assigned a ground potential ($V = 0$), whereas the opposing electrode was excited with an AC voltage ranging from 0 to 500 $V_{peak}$ at 8 kHz. In COMSOL, the frequency-domain solver uses the peak voltage amplitude as the excitation input. Electrically insulating boundary conditions were applied to all PDMS and channel-wall boundaries. The electrical conductivity of the fluid domain was set to 2 mS m$^{-1}$ to match the DEP buffer used experimentally. The resulting electric field distributions were subsequently used to evaluate the electric field strength and the electric field gradient, within the DEP region.

**Table S1** Outlet flow-rate calculations obtained from COMSOL line integrations of outlet velocity profiles under applied sample and sheath inlet flow rates of 5 and 10 $\mu$L min$^{-1}$, respectively. A channel height of $h = 120$ $\mu$m was used, corresponding to the fabricated microchannel height.

| **Outlet** | **Outlet width (Cut-line length)** | **Line integration formula** | **COMSOL expression** | **Flow rate ($\mu$l/min)** |
|---|---|---|---|---|
| CTC outlet-up | 700 $\mu$m | $Q_i = h \int_{\Gamma_i} u_n \, d\Gamma$ | spf.U*channelheight | 7.04 |
| Waste outlet | 150 $\mu$m | $Q_i = h \int_{\Gamma_i} u_n \, d\Gamma$ | spf.U*channelheight | 0.82 |
| CTC outlet-bottom | 700 $\mu$m | $Q_i = h \int_{\Gamma_i} u_n \, d\Gamma$ | spf.U*channelheight | 7.04 |

The imposed total inlet flow rate in the simulations was 15.00 $\mu$L min$^{-1}$, corresponding to the combined sample and sheath inlet flow rates. COMSOL line integrations yielded a total outlet flow rate of 14.90 $\mu$L min$^{-1}$ with a relative error of approximately 0.67%. The close agreement between the imposed inlet and calculated outlet flow rates confirms mass conservation throughout the computational domain and validates the COMSOL simulations used to compute outlet-specific flow rates and cell-fluxes.

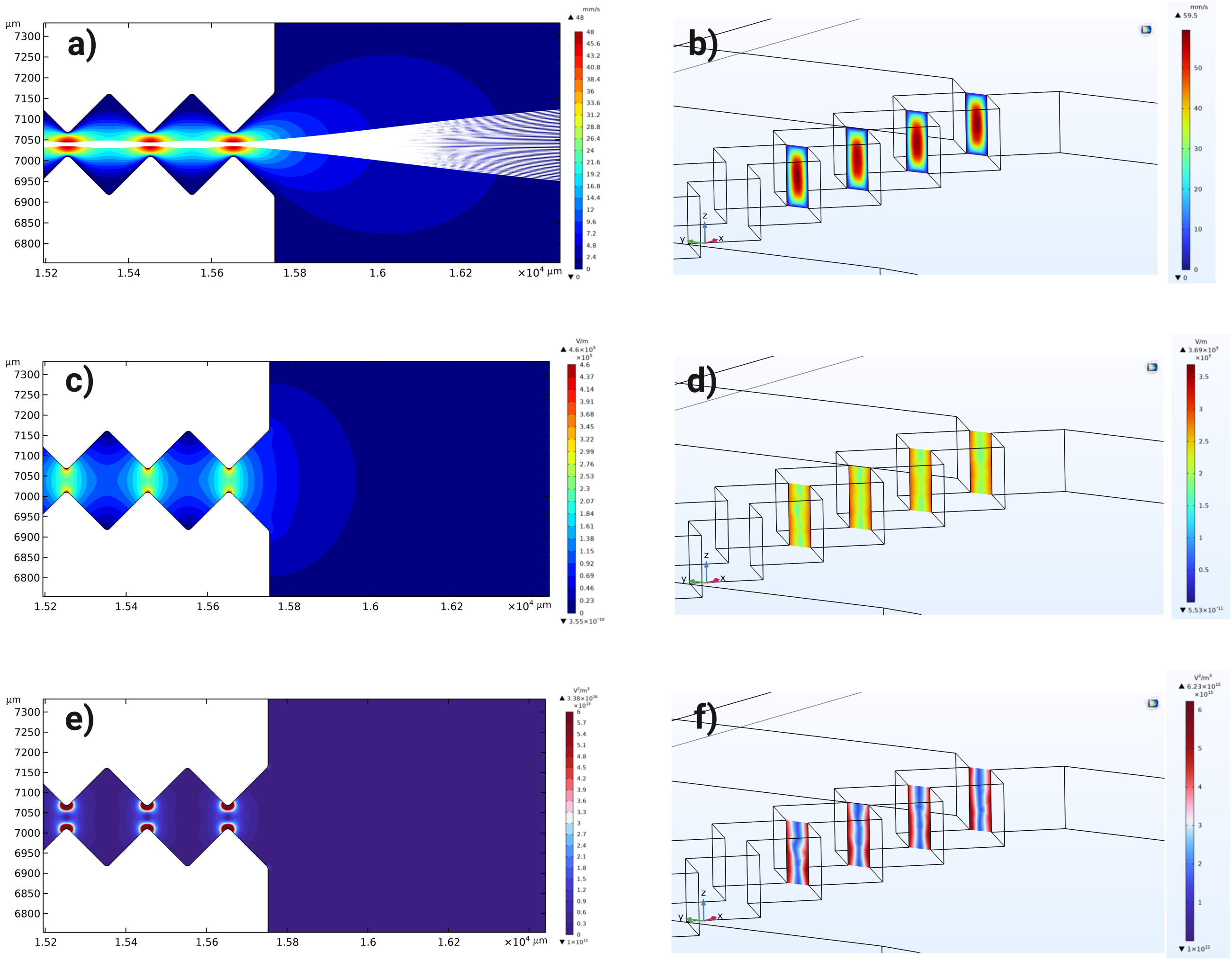


**Fig. S2** Finite-element simulation results for the final microfluidic device under the experimental operating conditions. (a) Two-dimensional flow velocity distribution demonstrating stable hydrodynamic focusing generated by sample and sheath flow rates of 5 and 10 $\mu$L min$^{-1}$, respectively. (b) Three-dimensional vertical cross-sectional view of the flow velocity profile showing the velocity distribution throughout the 120 $\mu$m channel height. (c) Two-dimensional electric field strength distribution within the DEP region at an applied voltage of 300 $V_{peak}$ (210 $V_{RMS}$) and 8 kHz. (d) Three-dimensional vertical cross-sectional view of the electric field strength demonstrating a nearly uniform electric field distribution throughout the channel height, except in the immediate vicinity of the insulating obstacle surfaces. (e) Two-dimensional distribution of the electric field gradient, $\nabla(E^2)$, relevant to dielectrophoretic force generation. (f) Three-dimensional vertical cross-sectional view of $\nabla(E^2)$ showing that the simulated electric field gradients exceed the reported threshold of approximately $10^{12}$ $V^2\,m^{-3}$ associated with significant dielectrophoretic manipulation of biological cells[1], confirming that the selected operating conditions generate electric field gradients sufficient for manipulation of cancer cells throughout the DEP region.

### S2. Fluorescent cell counting workflow used for multiple images and image stacks in ImageJ/Fiji

1. Open or import the experiment file (`.czi`), which represents an image stack, into ImageJ using Fiji.
   - In the import options, change the color mode to `Grayscale`. Cells may be stained with different fluorescent dyes (e.g., green, red, blue).
2. Convert the image stack type to 8-bit (`Image` → `Type` → `8-bit`).
3. Adjust the threshold of the image stack (`Image` → `Adjust` → `Threshold`).
   - Ensure the `Dark Background` option is checked. Set the method to `Default` and select the `Red` overlay.
   - Adjust the threshold bar until all fluorescently stained cells are fully included within the threshold range (0–255) while keeping the background as dark as possible. Note that the threshold value may vary between datasets due to differences in fluorescence intensity.
   - Click `Apply`. In the `Convert Stack to Binary` window, select the `Default` method and ensure the `Dark Background` option is enabled.
   - If the option `Calculate threshold for each image` is checked, uncheck it. Click `OK` and close the threshold window.
4. Add the region of interest (ROI).
   - Go to `Analyze` → `Tools` → `ROI Manager`.
   - In the ROI Manager window, select `More` → `Specify`. Define the rectangle width and height, and adjust the $x, y$ coordinates.
   - Check the option `Scaled units (microns)` and leave other options unchecked. Click `OK`.
   - Enable `Show All` in the ROI Manager window.
   - To display the ROI in every image of the stack, select `More` → `Options` and uncheck `Associate "Show All" ROIs with slices`.
   - Ensure that the ROI is selected in the ROI Manager before proceeding.
5. Click `Analyze` → `Measure` to record parameters such as area and mean value. Close the measurement window afterward without saving.
6. Perform particle analysis by selecting `Analyze` → `Analyze Particles`.
   - Set the particle size range (minimum and maximum) based on the expected projected cell area in the acquired images. For the cell lines used in this study, a range of 10–10,000 $px^2$ was found to provide accurate cell detection while excluding most debris.
   - Set the circularity to 0.10–1.00. The circularity threshold may be adjusted after visual inspection of the segmentation results to ensure that individual cells are accurately identified while excluding debris, fragmented cells, and cell aggregates.
   - Select `Outlines` in the `Show` option.
   - Enable `Summarize` and optionally `Exclude on Edges`. Leave other options unchecked and process all images in the stack.
7. The `Summary` window displays parameters such as image index, particle count, total area, and average size for each frame. The total number of counted cells can be calculated manually or obtained directly from the final frame of the image stack.
